# Dissecting adaptive traits with nested association mapping: Genetic architecture of inflorescence morphology in sorghum

**DOI:** 10.1101/748681

**Authors:** Marcus O. Olatoye, Sandeep R. Marla, Zhenbin Hu, Sophie Bouchet, Ramasamy Perumal, Geoffrey P. Morris

## Abstract

In the cereal crop sorghum (*Sorghum bicolor*) inflorescence morphology variation underlies yield variation and confers adaptation across precipitation gradients, but its genetic basis is poorly understood. Here we characterized the genetic architecture of sorghum inflorescence morphology using a global nested association mapping (NAM) population (2200 recombinant inbred lines) and 198,000 phenotypic observations from multi-environment trials for four inflorescence morphology traits (upper branch length, lower branch length, rachis length, and rachis diameter). Trait correlations suggest that lower and upper branch length are under largely independent genetic control, while lower branch length and rachis diameter are pleiotropic. Joint linkage and genome-wide association mapping revealed an oligogenic architecture with 1–22 QTL per trait, each explaining 0.1%–5.0% of variation across the entire NAM population. Overall, there is a significant enrichment (2.4-fold) of QTL colocalizing with homologs of grass inflorescence genes, notably with orthologs of maize (*Ramosa2*) and rice (*Aberrant Panicle Organization1, TAWAWA1*) inflorescence regulators. In global georeferenced germplasm, allelic variation at the major inflorescence morphology QTL is significantly associated with precipitation gradients, consistent with a role for these QTL in adaptation to agroclimatic zones. The findings suggest that global inflorescence diversity in sorghum is largely controlled by oligogenic, epistatic, and pleiotropic variation in ancestral regulatory networks. This genotype-phenotype trait dissection in global germplasm provides a basis for genomics-enabled breeding of locally-adapted inflorescence morphology.

## INTRODUCTION

Understanding the genetic architecture of complex traits in crops provides insights into crop evolution and guidance on breeding strategies. Adaptive traits are phenotypic characteristics that are subject to selection towards an optimum for a particular environment (Barrett and Hoekstra, 2011). Genetic architecture describes the structure of the genotype-phenotype map for complex traits in populations: the number of loci, distribution of effect size, frequencies of alleles, gene action (dominance and epistasis), and the degree of linkage and pleiotropy (Holland, 2007). Trait variation in a population may shift from oligogenic to polygenic architecture as a population moves towards an optimum (i.e. Fisher’s geometric model) (Orr, 2005; Tenaillon, 2014). Thus, characterizing genetic architecture of complex traits is key step to bridge theoretical understanding (e.g. evolutionary, metabolic, or developmental drivers) and applied outcomes (e.g. crop and livestock breeding strategies, or management of human genetic disorders) (Hansen, 2006; Timpson *et al.*, 2018). For instance, molecular breeding strategies are guided by genetic architecture, with marker-assisted backcross for monogenic traits, marker-assisted recurrent selection for oligogenic traits, and genomic selection for polygenic traits (Bernardo, 2008).

Divergence of adaptive traits often results in genetic differentiation and population structure that hinders effective characterization of their genetic architecture (Myles *et al.*, 2009; Brachi *et al.*, 2011). Genome-wide association studies (GWAS) in diverse natural populations have been widely used to characterize genetic architecture but are limited by a fundamental tradeoff when causative variants (i.e. the oligogenic component) are confounded with polygenic variation. Models without population and/or kinship terms partition the colinear variance into the monogenic/oligogenic term (leading to false positive associations) while models with population and/or kinship terms partition colinear variation into polygenic terms (leading to false negatives) (Bergelson and Roux, 2010). In a nested association mapping (NAM) population, controlled crosses between the common parent and diverse founders breaks up population structure, increasing power for QTL detection (Myles *et al.*, 2009). In addition, the larger population size in most NAM populations mitigates the Beavis effect, the overestimation of QTL effect size that occurs in small populations (Utz *et al.*, 2000). NAM has greatly facilitated the characterization of genetic architecture in species where controlled crosses are feasible, including many major crops (Buckler *et al.*, 2009; Maurer *et al.*, 2015; Bajgain *et al.*, 2016; Bouchet *et al.*, 2017).

Inflorescence morphology is a key component of crop adaptation and yield (Harlan and de Wet, 1972; Cooper *et al.*, 2014). Homologous variation of inflorescence among cereals has long been noted (Vavilov, 1922) and inflorescence morphology has been a valuable system to investigate the evolutionary dynamics and molecular basis of genetic architecture in plants (Hermann and Kuhlemeier, 2011; Zhang and Yuan, 2014). Analysis of inflorescence mutants has revealed regulatory networks with genes controlling hormonal biosynthesis, hormone transport, signal transduction, and transcriptional regulation (Zhang and Yuan, 2014). Comparative studies indicate that components of inflorescence regulatory networks are largely conserved across grass species, but that substantial variation in ancestral regulatory networks exists within and among species (Kellogg, 2007; Barazesh and McSteen, 2008; Tanaka *et al.*, 2013; Huang *et al.*, 2017). However, since most inflorescence regulators were identified via mutant screens, the role of these ancestral genes in natural variation or adaptive divergence of inflorescence morphology is not well understood (Brown *et al.*, 2011; Crowell *et al.*, 2016; Wu *et al.*, 2016). In addition, studies of natural variation may reveal genes not yet identified via mutant analysis.

Sorghum is a source of food, feed, and bioenergy in many parts of the world, especially important to smallholder farmers in semi-arid regions (National Research Council, 1996). Sorghum has diffused to contrasting agroclimatic zones, and harbors abundant variation in traits such as height, leaf architecture, and inflorescence morphology. Variation in inflorescence morphology underlies yield components (Brown *et al.*, 2006; Witt Hmon *et al.*, 2013) and local adaptation to agroclimatic zones defined by precipitation gradients (De Wet *et al.*, 1972; Olatoye *et al.*, 2018). The five major botanical races of sorghum are largely defined based on inflorescence morphology (Harlan and de Wet, 1972). For instance, guinea sorghums with long open panicles predominate in humid zones while durra sorghums with short compact panicles predominate in arid zones (Lasky *et al.*, 2015; Olatoye *et al.*, 2018). However the genetic architecture of inflorescence morphology in sorghum is poorly understood, and no genes have been characterized (Brown *et al.*, 2008; Witt Hmon *et al.*, 2013). In this study we used nested association mapping to characterize the genetic architecture of inflorescence morphology variation in global sorghum. Our results reveal that global sorghum inflorescence variation is under the control of oligogenic, epistatic, and pleiotropic loci, consistent with Fisher’s geometric model under disruptive selection.

## MATERIALS AND METHODS

### Plant materials and phenotyping

The sorghum NAM population was derived from a cross between an elite U.S. common parent RTx430 and 10 diverse founders that originated from different agroclimatic zones, thereby capturing a wide genetic and morphological diversity (Supplementary Table 1, Supplementary Figure 1) (Bouchet *et al.*, 2017). Each diverse parent and its RILs represent a family of 200–233 RILs making a total of 2220 RILs in the population. Field phenotyping experiments were conducted in semi-arid (Hays, Kansas; 38.86, −99.33) and humid-continental (Manhattan, Kansas; 39.21, −96.59) environments for two years (2014 and 2015) to represent a typical range of growing conditions. Each site-by-year was regarded as one environment (Table 1). In the second year (2015), RILs were randomized within maturity blocks of families in a row-column design based the first-year flowering data. Each row (corresponding to a plot) was 3 m with 1 m alleys between ranges.

The NAM RILs were phenotyped at F6:7 and F6:8 generations for upper primary branch length (UBL), lower primary branch length (LBL), rachis length (RL), and rachis diameter (RD) (Supplementary Figure 2). Three random panicles were collected from each plot after physiological maturity and subsequently used for phenotyping. Inflorescence morphology traits were measured using barcode rulers (1 mm precision) and barcode readers (Motorola Symbol CS3000 Series Scanner, Chicago IL, USA). RL was measured as the distance from apex of the panicle to the point of attachment of the lowest rachis lower primary branch (Brown *et al.*, 2006). RD was measured using a digital Vernier caliper as the diameter of the peduncle at the point of attachment of the bottommost rachis lower primary branch. For UBL, three primary branches were randomly detached from the apex of the panicle. For LBL, three primary branches were randomly detached from the region closest to the peduncle for two panicles (Supplementary Figure 2).

### Genomic data analysis

Genotyping-by-sequencing of the NAM population and diverse global germplasm was previously described (Bouchet *et al.*, 2017; Hu *et al.*, 2019). Briefly, Illumina sequence reads were aligned to the BTx623 reference genome version 3 using Burrow Wheeler Aligner 4.0 and SNP calling was done using TASSEL-GBS 5.0 (Glaubitz *et al.*, 2014). For the current study, missing data imputation was done in two stages using Beagle 4 (Browning and Browning, 2013). The NAM population and the sorghum association mapping population (SAP) GBS data were first extracted from the build. Filtering was conducted to remove markers with (i) tri-allelic SNPs, (ii) missing data in more than 80% of individuals, or (iii) < 3% minor allele frequency prior to imputation. The NAM population and SAP were imputed separately and each germplasm set was filtered for MAF > 0.05. NAM RILs with >10% heterozygosity were dropped from the analysis.

### Phenotype and heritability analysis

Phenotypic data analysis was carried out using R programming language and SAS (SAS Institute Inc., Cary, NC, USA). All traits were tested for normality and the only trait (UBL) with significantly skewed distribution was log transformed. Analysis of variance was performed for each trait using *aov* function in R. The best linear unbiased prediction (BLUP) of each trait was estimated using data from five environments with *lmer* function in *LME4* package in R (Bates *et al.*, 2014) with genotype, environment, and genotype-environment interactions fitted as random effects (Wu *et al.*, 2016). The variance components used for broad sense heritability (*H*^2^) were estimated using the maximum likelihood method by PROC VARCOMP of the SAS software (SAS Institute Inc., Cary, NC, USA). RIL-nested-within-family and RIL-nested-within-family by environment interaction were fit as random effects. The resulting variance components were used to estimate the broad sense heritability (*H*^2^) following equation 1 in (Hung *et al.*, 2012) as:

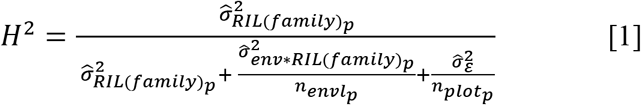

where 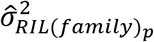 is the variance component of RILs nested within family *p, n_envl_p__* is the harmonic mean of the number of environments in which each RIL was observed, and *n_plot_p__* the harmonic mean of the total number of plots in which each RIL. Pearson pairwise correlation between traits was estimated using the residuals derived from fitting a linear model for family and trait phenotypic means:

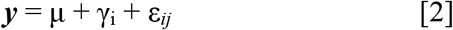

where y is the vector of phenotypic data, μ is the overall mean, γ_i_ is the term for the NAM families, and ε_*ij*_ is the residual.

### Joint linkage mapping

Joint linkage analysis was performed using 92,391 markers and 2220 RILs. This approach is based on forward inclusion and backward elimination stepwise regression approaches implemented in TASSEL 5.0 stepwise plugin (Glaubitz *et al.*, 2014). The family effect was accounted for as a co-factor in the analysis. First, a nested joint linkage (NJL) model was fitted where markers were nested within families (Poland *et al.*, 2011; Würschum *et al.*, 2012). In addition, a non-nested joint linkage model (JL), where markers were not nested within families, was used due to its higher predictive power than NJL (Würschum *et al.*, 2012). Entry and exit F_test_ values were set to 0.001 and based on 1000 permutations, the *P*-value threshold was set to 1.84 × 10^−6^. The JL model was specified as:

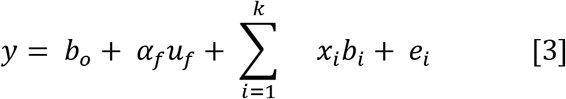

where *b*_0_ is the intercept, *u_f_* is the effect of the family of founder *f* obtained in the cross with the common parent (RTx430), α_*f*_ is the coefficient matrix relating *u_f_* to *y, b_i_* is the effect of the *i*^th^ identified locus in the model, *x_i_* is the incidence vector that relates *b_i_* to *y* and *k* is the number of significant QTL in the final model (Yu *et al.*, 2008).

### Genome-wide association studies

GWAS was performed for all traits using 92,391 markers and 2220 RILs using BLUPs adjusted by environments. The multi-locus-mixed model (MLMM) approach (Segura *et al.*, 2012) implemented in R was used for GWAS in the NAM population, as described previously (Bouchet *et al.*, 2017). The MLMM approach performs stepwise regression involving both forward and backward regressions, accounts for major loci and reduces the effect of allelic heterogeneity. The family effect was fitted as a co-factor and a random polygenic term (kinship relationship matrix) was also accounted for in the MLMM model. Bonferroni correction with α = 0.05 was used to determine the cut-off threshold for each trait association (α/total number of markers = 5.4 × 10^−7^).

For comparison with NAM, GWAS was performed in the sorghum association panel (SAP; 334 accessions) (Casa *et al.*, 2008) using general linear model (GLM) and compressed mixed linear model (CMLM) with the GAPIT R package (Lipka *et al.*, 2012) to match a previous study (Morris *et al.*, 2013). The GLM (naive model) did not account for population structure and was specified as:

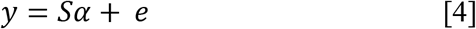

where **y** is the vector of phenotypes, α is a vector of SNPs effects, and *e* is the vector of residual effects, and **S** is the incident matrix of 1s and 0s relating *y* to α. The CMLM model (full model) accounted for population structure and polygenic background effects (kinship) was specified as:

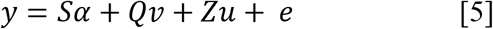

where **y** is the vector of phenotype, and *u* is a vector of random genetic background effects. **X, Q**, and **Z** are incident matrices of 1s and 0s relating **y** to β and *u* (Yu *et al.*, 2006). The phenotypic data in the SAP used for GWAS is from a previous study (Brown *et al.*, 2008; Morris *et al.*, 2013). A custom script was used to identify QTL that overlapped within a 50 kb window between the NAM and GWAS (GLM or CMLM) mapping results for LBL and RL.

### Effect size and allele frequency estimation

Allele frequencies at the SNPs were calculated using snpStats package in R (Clayton 2015). The additive effect size of QTL within and across families were estimated as the difference between the mean of the two homozygous classes for each QTL divided by two. The additive effect of each QTL was estimated relative to RTx430. The sum of squares of QTLs divided by the total sum of squares gave the proportion of variance explained. To estimate within-family variation explained by each QTL, a regression model was fit with terms for family and QTL nested within family as fixed effects (Würschum *et al.*, 2012):

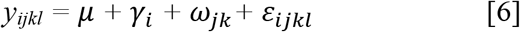

where *y_ijkl_* is the phenotype, γ_i_ is the family term, ω_*jk*_ is the term for QTL nested within family, and ε_*ijk*_ is the residual.

### Grass homologs search around identified loci and enrichment analysis

A set of known inflorescence genes (n = 29) that control inflorescence morphology in grasses were compiled from literature consisting of 17 maize genes, eight rice genes, three sorghum genes, and one foxtail millet gene (Supplementary File 1). Based on this candidate gene set, 135 sorghum homologs were found using Phytozome (Goodstein *et al.*, 2012). A custom R script was used to search for homologs within 150 kb window upstream and downstream of each association, based on the LD decay rate in the NAM population (Hu *et al.*, 2019). The enrichment of *a priori* genes around identified QTL was performed using chi square test to compare *a priori* gene colocalization around identified QTL with colocalization of QTL with random genes from the sorghum genome version 3.1 gff3 file on Phytozome.

### Geographic allele distribution and climatic association of inflorescence QTL

The distribution of the alleles of QTL markers underlying inflorescence variation was visualized on a global geographic map with national boundaries. Markers of QTL associated with inflorescence morphology that also colocalized with sorghum orthologs of maize and rice inflorescence genes were first selected from the genetic data of a global georeferenced sorghum germplasm (number of accession = 2227, number of SNP markers = 431691). The alleles of the NAM markers found in the georeferenced germplasm were then plotted based on the geographic coordinates of the individuals in which they are present. Furthermore, climatic genome wide association was performed between the annual mean precipitation and the genetic data of the georeferenced sorghum global germplasm using both the naive model (GLM) and the mixed model that accounted for kinship only.

### Data availability

Phenotype and genotype data are available at FigShare: *[add after acceptance].* File S1 contains detailed descriptions of QTL information, *a priori* gene list and *a priori* genes that colocalized with QTL. File S2 contains heatmap of QTL effects within NAM families. File S3 contains detailed description of associations that colocalized between NAM, GLM, and CMLM and results of association of inflorescence QTL markers with precipitation for both GLM and MLM. The NAM population seeds are available from the USDA National Plant Germplasm System (https://www.ars-grin.gov/). Raw sequencing data for the NAM population are published (Bouchet *et al.*, 2017) and available in the NCBI Sequence Read Archive under project accession SRP095629 and on Dryad Digital Repository (doi:10.5061/dryad.gm073).

## RESULTS

### Variation of inflorescence morphology in the NAM population

Phenotypic measurements were collected for four inflorescence morphology traits across five environments, representing over 198,000 observations. The number of RILs in each family ranged from 202 in the Segaolane family to 233 in the SC265 family (Supplementary Table 1). Significant genotypic differences were observed for all four inflorescence traits (Table 2). The broad-sense heritability estimates for all four traits were high, ranging from 0.59 to 0.92. The SC265 and SC283 families had the longest lower branches (mean across RILs of 99 mm). The SC283 family had the longest upper branches (mean across RILs of 64 mm). The SC265 and Segaolane families had the longest rachis, with mean lengths of 316 mm and 305 mm, respectively. The largest rachis diameters were observed in the Ajabsido, Macia, and SC35 families (a mean of ~9.5 mm across RILs in each family). Phenotypic variation distribution within families showed that in some families the mean trait value of the RILs was greater than the mean of either parent (Figure 1). The highest trait-by-trait phenotypic correlations were for RL and LBL (*r* = 0.71; *P*-value < 0.01). By contrast, UBL and LBL had a low positive correlation (*r* = 0.19; *P*-value < 0.01), and RL had no correlation with either UBL or RD (Figure 2).

**Figure 1:**
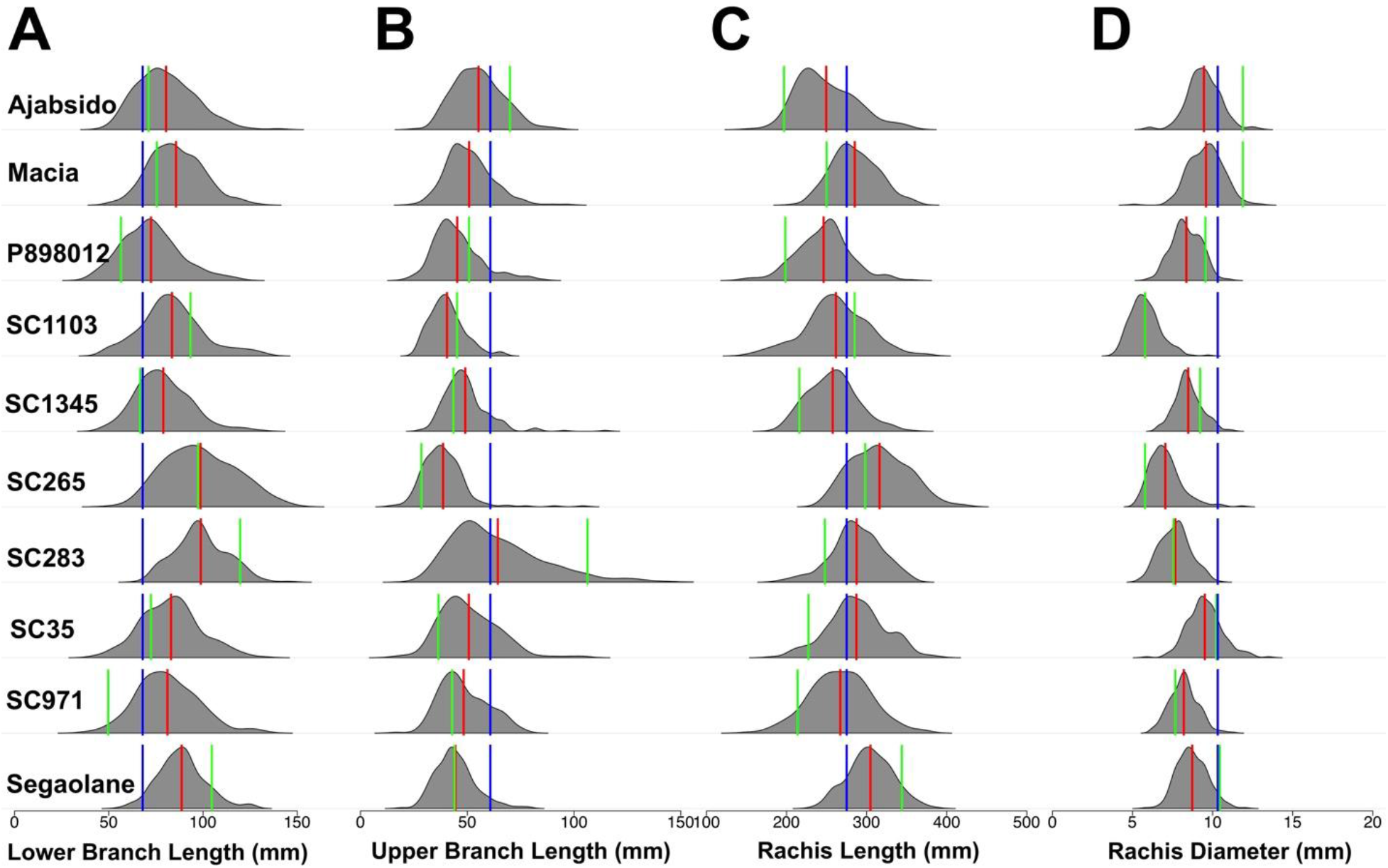
Phenotypic distribution of sorghum inflorescence morphology. Phenotypic distribution of line means for inflorescence traits (A) lower branch length, (B) upper branch length, (C) rachis length, and (D) rachis diameter. Blue lines indicate trait value for the common parent (RTx430), green lines indicate mean trait values for each of the other parents (listed on the left), and red lines indicate the trait mean across each recombinant inbred line family.

**Figure 2:**
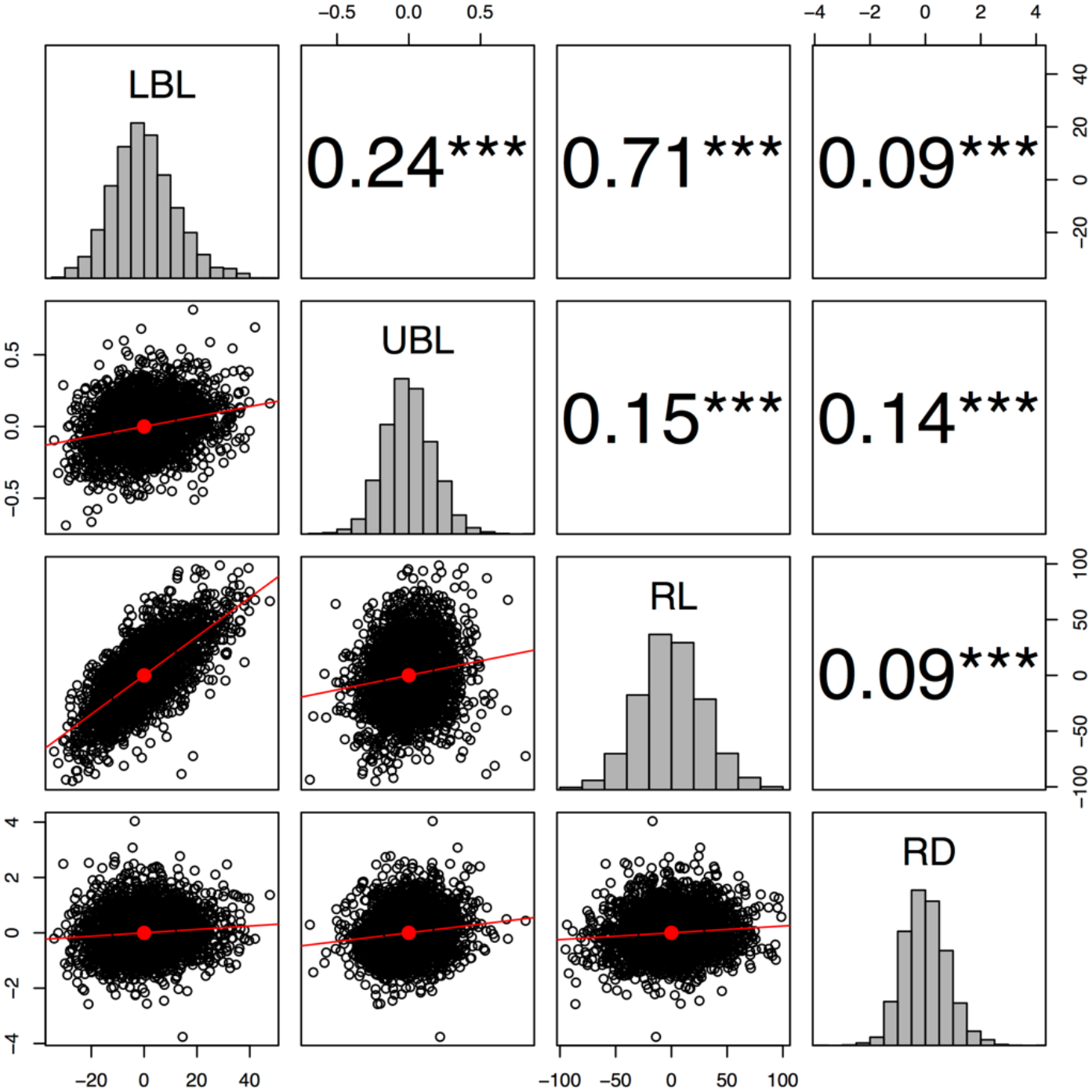
Pairwise correlation among inflorescence morphology traits. Pearson correlation between residuals of the regression of the family on the the best linear unbiased predictors (BLUPs) of lower branch length (LBL), upper branch length (UBL), rachis length (RL), and rachis diameter (RD) significant at 0.05, 0.01 and 0.001 (*, **, and ***). BLUPs were estimated across five environments (year-by-location).

**Table 1:**
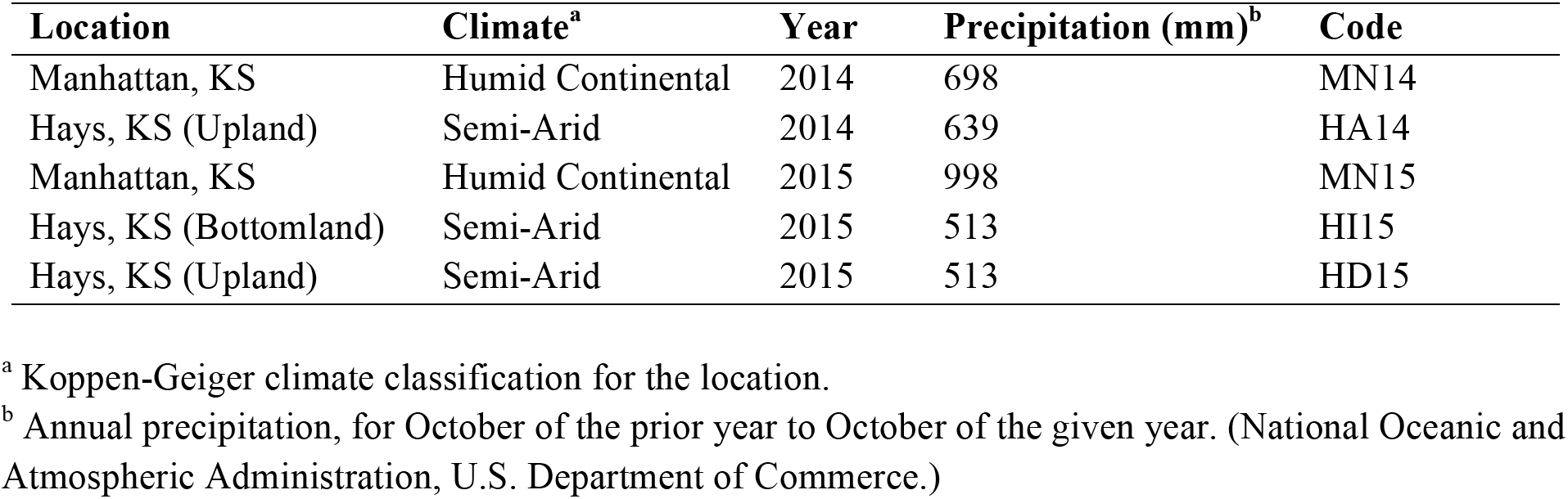
Summary of field experiments using the nested association mapping population.

**Table 2:** Mean, range, and broad sense heritability (*H*^2^) for lower branch length (LBL), upper branch length (UBL), rachis length (RL), and rachis diameter (RD).

| **Trait<sup>a</sup>** | **Range (mm)** | **Mean (mm)** | **$H^2$** |
| --- | --- | --- | --- |
| LBL*** | 267 – 176 | 82 | 0.86 |
| UBL* | 7 – 170 | 48 | 0.85 |
| RL*** | 111 – 465 | 274 | 0.92 |
| RD*** | 3.8 – 13.5 | 8.3 | 0.59 |

<sup>a</sup> Significant genotypic differences given by \*, \*\*, \*\*\* at 0.05, 0.01 and 0.001, respectively.

### QTL variation in the NAM population

A total of 116,405 SNPs were obtained after SNP calling, imputation, and filtering (minimum MAF = 5%). After filtering for 0.96 inbreeding coefficient, a total of 92,391 markers were identified. Significant QTL associations were observed for all traits when using MLMM, JL, and NJL models (Figure 3, Supplementary Figure 3–4). MLMM identified nine significant associations in total for all traits. The JL model identified 81 QTL, while the NJL model identified 40 QTL across all traits (Supplementary File 1 and Supplementary Table 2). Allele frequencies at the QTL ranged from 0.05 to 0.48 (Supplementary File 1). The proportion of within-family variation explained by all QTL (i.e. an estimate of the oligogenic component) varied substantially among traits, with 12%, 37%, 31%, and 21% of variation explained by QTL for UBL, LBL, RL, and RD, respectively. Within-family and across-family effect of each QTL for NJL and JL models were estimated relative to RTx430 (Supplementary File 2). LBL QTL qSbLBL7.5960 explained the largest proportion of variation among all QTL identified in this study (Table 3).

**Figure 3:**
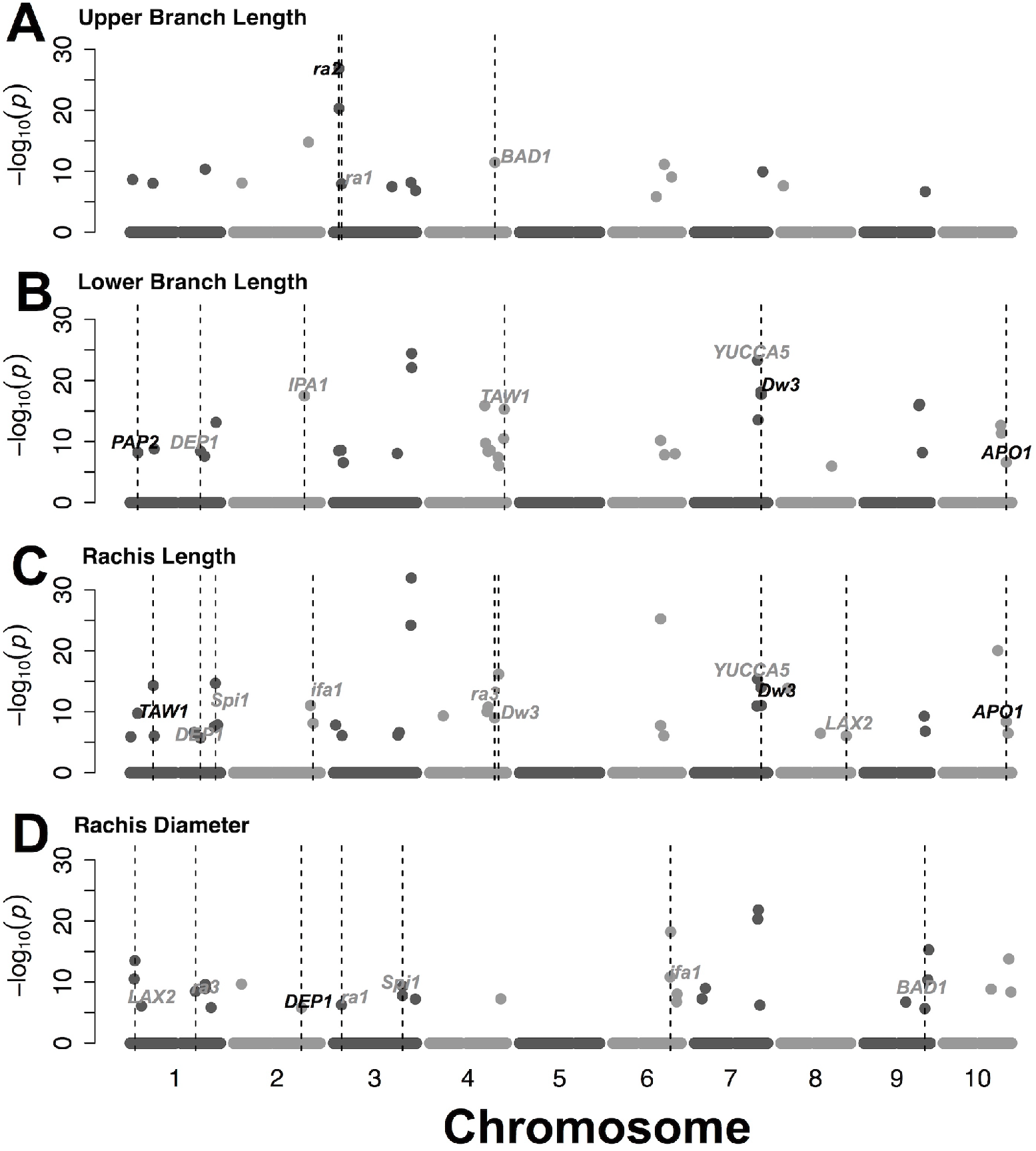
QTL mapping for inflorescence morphology using joint linkage model. Genome positions of loci associated with (A) upper branch length, (B) lower branch length, (C) rachis length, and (D) rachis diameter. *A priori* candidate genes that colocalize with QTL within 150 kb are noted as follows. Black text indicates putative sorghum orthologs of *a priori* candidate genes while gray text indicates paralogs.

**Table 3:** Inflorescence morphology QTL that explain > 1.5% of variation across the NAM population.

| QTL Code | MAF<sup>a</sup> | PVE<sup>b</sup> | Trait<sup>c</sup> | Gene Colocalization<sup>d</sup> |
| --- | --- | --- | --- | --- |
| qSbUBL3.475 | 0.18 | 3.8 | UBL | P |
| qSbUBL3.476 | 0.26 | 3.2 | UBL | P |
| qSbUBL2.672 | 0.43 | 2.1 | UBL | A |
| qSbUBL3.734 | 0.23 | 1.8 | UBL | A |
| qSbUBL6.461 | 0.11 | 1.7 | UBL | A |
| qSbUBL3.524 | 0.05 | 1.6 | UBL | A |
| qSbRL10.488 | 0.39 | 2.6 | RL | A |
| qSbRL3.699 | 0.32 | 2.5 | RL | A |
| qSbRL6.428 | 0.23 | 2.2 | RL | A |
| qSbRL4.514 | 0.16 | 1.8 | RL | P |
| qSbRL1.785 | 0.36 | 1.7 | RL | A |
| qSbRL1.216 | 0.38 | 1.5 | RL | A |
| qSbRL3.694 | 0.44 | 1.5 | RL | A |
| qSbRL6.428 | 0.32 | 1.5 | RL | A |
| qSbLBL7.598 | 0.18 | 4.4 | LBL | P |
| qSbRL7.598 | 0.18 | 3.0 | RL | P |
| qSbLBL7.596 | 0.18 | 4.6 | LBL | A |
| qSbLBL7.566 | 0.23 | 4.3 | LBL | A |
| qSbLBL7.569 | 0.18 | 3.6 | LBL | A |
| qSbLBL4.524 | 0.1 | 3.1 | LBL | A |
| qSbLBL4.493 | 0.41 | 2.8 | LBL | A |
| qSbLBL4.669 | 0.05 | 2.6 | LBL | A |
| qSbLBL4.621 | 0.17 | 2.5 | LBL | A |
| qSbLBL4.501 | 0.47 | 2.3 | LBL | A |
| qSbLBL2.636 | 0.26 | 2.2 | LBL | A |
| qSbLBL10.519 | 0.13 | 1.8 | LBL | A |
| qSbLBL2.635 | 0.43 | 1.7 | LBL | P |
| qSbLBL4.542 | 0.3 | 1.7 | LBL | A |
| qSbLBL3.702 | 0.37 | 1.6 | LBL | A |
| qSbLBL7.571 | 0.34 | 1.6 | LBL | A |
| qSbLBL9.493 | 0.38 | 1.6 | LBL | A |
| qSbLBL3.702 | 0.37 | 1.5 | LBL | A |

<sup>a</sup> MAF: Minor allele frequency.
<sup>b</sup> PVE: Proportion of variation explained.
<sup>c</sup> Lower branch length (LBL), upper branch length (UBL), rachis length (RL), and rachis diameter (RD).
<sup>d</sup> “P” (present) or “A” (absence) of colocalization with *a priori* candidate gene (within 150 kb from QTL).

### QTL colocalization and enrichment with *a priori* candidate genes

To assess the overall role of variation at ancestral inflorescence regulators, we performed colocalization and enrichment analysis between the QTL and a set of *a priori* candidate genes containing sorghum homologs of rice, maize, and foxtail millet genes (n = 135). NAM QTL were significantly enriched for colocalization with *a priori* candidate genes (2.4-fold enrichment; *P*-value < 0.001). Of 123 unique QTL, 28 colocalized with *a priori* genes. Among the QTL that overlapped with *a priori* candidate genes, two QTL were inside the gene, three QTL were <15 kb from the gene, 16 unique QTL were 15–100 kb from the gene, and eight unique QTL were 100–150 kb from the genes (Table 4 and Supplementary File 1). Overall, 24 genes colocalized with inflorescence QTL, while 111 *a priori* candidate genes (of 135) did not overlap with any inflorescence QTL (Supplementary File 1).

**Table 4:**
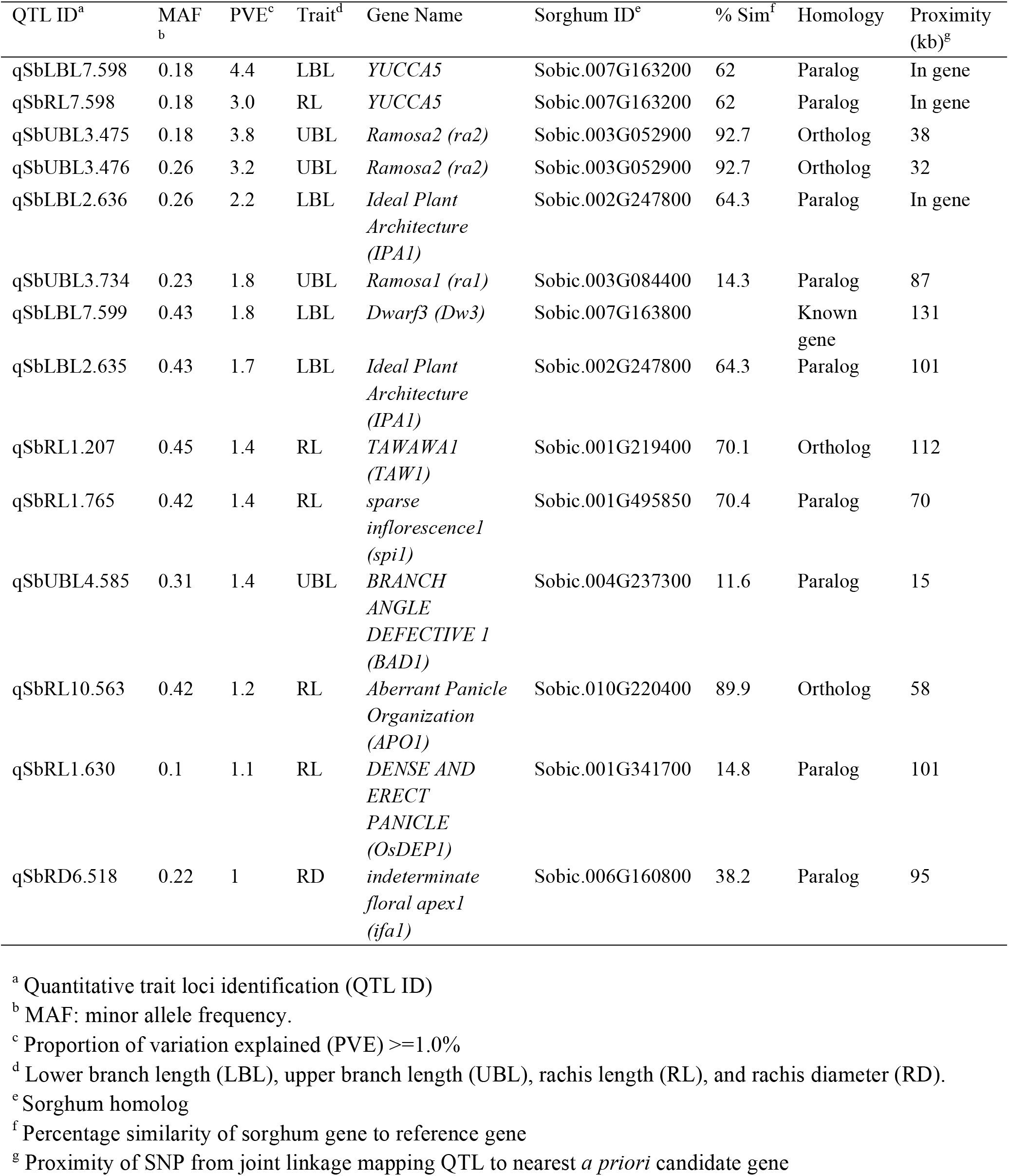
Joint linkage mapping QTL that colocalize with *a priori* candidate genes.

| QTL ID<sup>a</sup> | MAF<sup>b</sup> | PVE<sup>c</sup> | Trait<sup>d</sup> | Gene Name | Sorghum ID<sup>e</sup> | % Sim<sup>f</sup> | Homology | Proximity (kb)<sup>g</sup> |
| --- | --- | --- | --- | --- | --- | --- | --- | --- |
| qSbLBL7.598 | 0.18 | 4.4 | LBL | *YUCCA5* | Sobic.007G163200 | 62 | Paralog | In gene |
| qSbRL7.598 | 0.18 | 3.0 | RL | *YUCCA5* | Sobic.007G163200 | 62 | Paralog | In gene |
| qSbUBL3.475 | 0.18 | 3.8 | UBL | *Ramosa2 (ra2)* | Sobic.003G052900 | 92.7 | Ortholog | 38 |
| qSbUBL3.476 | 0.26 | 3.2 | UBL | *Ramosa2 (ra2)* | Sobic.003G052900 | 92.7 | Ortholog | 32 |
| qSbLBL2.636 | 0.26 | 2.2 | LBL | *Ideal Plant Architecture (IPA1)* | Sobic.002G247800 | 64.3 | Paralog | In gene |
| qSbUBL3.734 | 0.23 | 1.8 | UBL | *Ramosa1 (ra1)* | Sobic.003G084400 | 14.3 | Paralog | 87 |
| qSbLBL7.599 | 0.43 | 1.8 | LBL | *Dwarf3 (Dw3)* | Sobic.007G163800 |  | Known gene | 131 |
| qSbLBL2.635 | 0.43 | 1.7 | LBL | *Ideal Plant Architecture (IPA1)* | Sobic.002G247800 | 64.3 | Paralog | 101 |
| qSbRL1.207 | 0.45 | 1.4 | RL | *TAWAWA1 (TAW1)* | Sobic.001G219400 | 70.1 | Ortholog | 112 |
| qSbRL1.765 | 0.42 | 1.4 | RL | *sparse inflorescence1 (spi1)* | Sobic.001G495850 | 70.4 | Paralog | 70 |
| qSbUBL4.585 | 0.31 | 1.4 | UBL | *BRANCH ANGLE DEFECTIVE 1 (BAD1)* | Sobic.004G237300 | 11.6 | Paralog | 15 |
| qSbRL10.563 | 0.42 | 1.2 | RL | *Aberrant Panicle Organization (APO1)* | Sobic.010G220400 | 89.9 | Ortholog | 58 |
| qSbRL1.630 | 0.1 | 1.1 | RL | *DENSE AND ERECT PANICLE (OsDEP1)* | Sobic.001G341700 | 14.8 | Paralog | 101 |
| qSbRD6.518 | 0.22 | 1 | RD | *indeterminate floral apex1 (ifa1)* | Sobic.006G160800 | 38.2 | Paralog | 95 |

<sup>a</sup> Quantitative trait loci identification (QTL ID)
<sup>b</sup> MAF: minor allele frequency.
<sup>c</sup> Proportion of variation explained (PVE) $\geq 1.0\%$
<sup>d</sup> Lower branch length (LBL), upper branch length (UBL), rachis length (RL), and rachis diameter (RD).
<sup>e</sup> Sorghum homolog
<sup>f</sup> Percentage similarity of sorghum gene to reference gene
<sup>g</sup> Proximity of SNP from joint linkage mapping QTL to nearest *a priori* candidate gene

### Comparison of NAM and diversity panel GWAS

NAM provides an independent approach to validate GWAS QTL from diversity panels and assess the relative performance of GWAS models. We compared the inflorescence loci identified in the NAM with GWAS QTL for LBL and RL identified in the SAP, identifying colocalization (within 50 kb) between NAM QTL SNPs and top 5% SNP associations in the GLM or CMLM (Figure 4 and Supplemental File 2). For LBL, the comparison revealed 26 overlaps between NAM versus GLM, and 20 overlaps between NAM versus CMLM. For RL, the comparison revealed 17 overlaps for both NAM versus GLM and NAM versus CMLM. To identify gene candidate that are supported by multiple mapping approaches, *a priori* candidate genes were catalogued in overlapping NAM and GWAS QTL (Supplementary File 3). For LBL, five *a priori* candidate genes colocalized with overlapping NAM and GLM QTL, while two *a priori* candidate genes colocalized with overlapping NAM and CMLM QTL. Similarly for RL, six *a priori* candidate genes colocalized with overlapping NAM and GLM QTL, and six genes colocalized with overlapping NAM and CMLM QTL.

**Figure 4:**
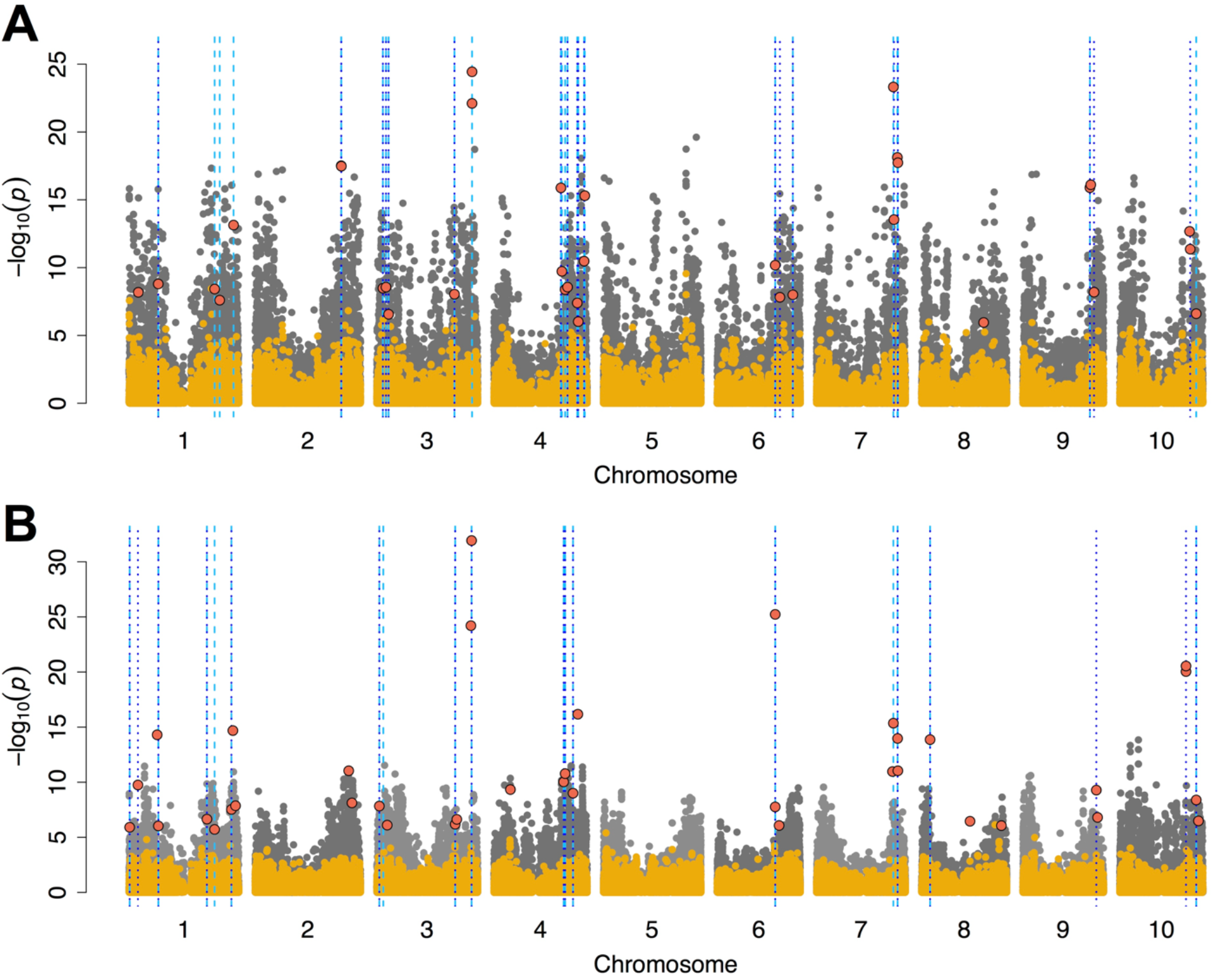
Comparison of joint linkage in a NAM population versus genome-wide association in a diversity panel. Manhattan plot for the comparison of genome-wide association approaches for (A) lower branch length and (B) rachis length using general linear model (GLM) in gray, compressed mixed linear model (CMLM) in yellow, and NAM joint linkage (JL) model in red. Broken lines in light blue and deep blue note colocalization between NAM and GLM (50 kb window), and between NAM and CMLM (50 kb window), respectively. GLM and CMLM were carried out in the sorghum association mapping panel (SAP, n = 334) and NAM (n = 2200).

### Geographic distribution of allele and environment-marker associations

A strong geographic pattern of distribution of SNP alleles associated with inflorescence morphology was observed (Figure 5). The LBL-associated C allele (S10_56303321) near the sorghum ortholog of *APO1* was predominant in Horn of Africa region, Yemen, Southern Africa, Southern India, and Eastern China. The T allele was predominant in West Africa. For the marker (S3_4750709) associated with upper branch length that colocalized with the sorghum ortholog of *ramosa2*, the allele was predominant in India, south-eastern Africa, and Sahelian region of West Africa. While the G allele was predominant in Nigeria and tropical West Africa, and eastern China. For the LBL-associated SNP (S7_59751994) near *YUCCA5* (i.e. the *sparse inflorescence1* paralog), one allele was predominant in West Africa and India, while the other allele was predominant in Southeastern Africa. All the QTL markers were found to be significantly associated with precipitation under the GLM but not with the MLM that accounted for kinship (File S3).

**Figure 5:**
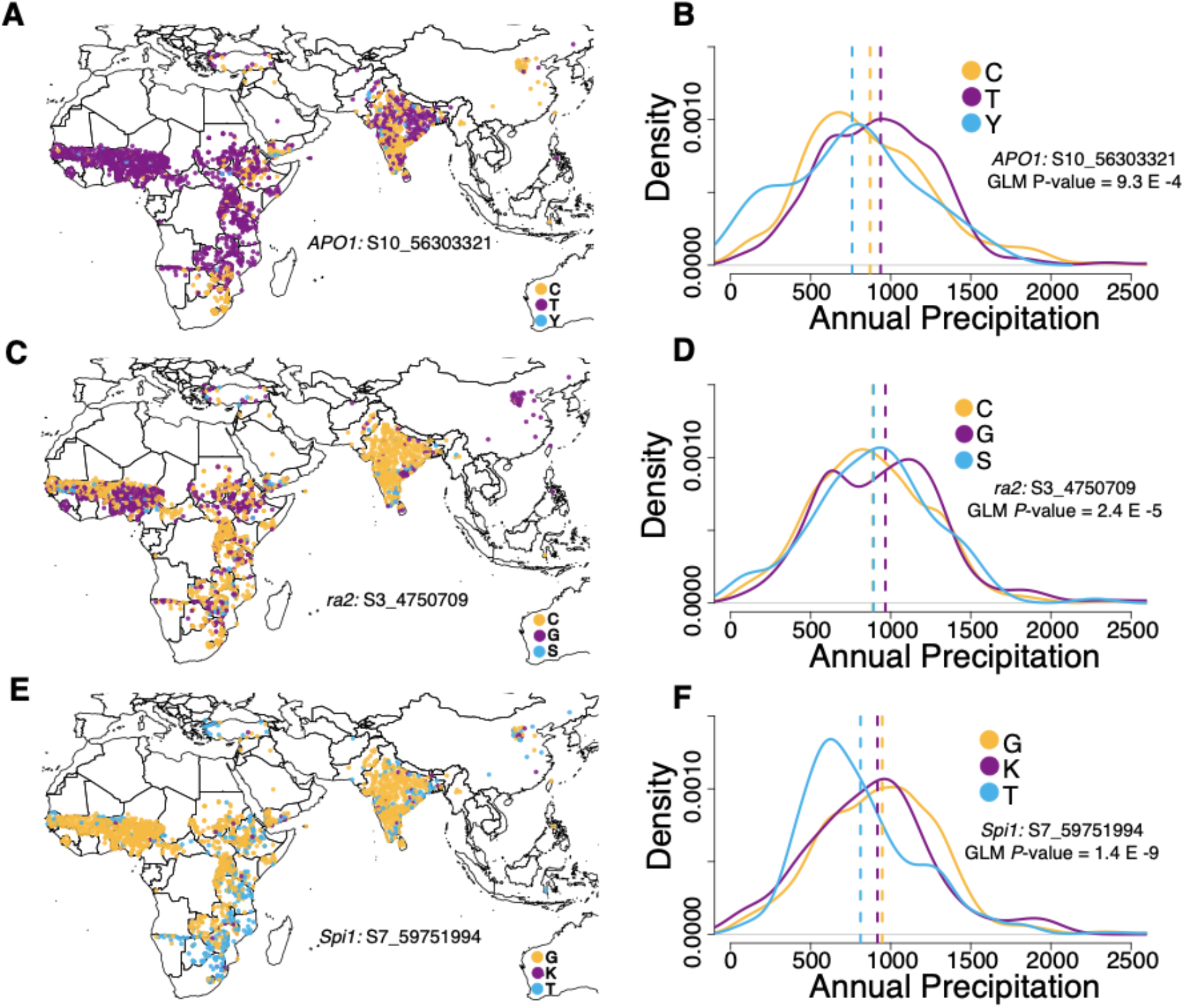
Global geographic allele distribution of inflorescence-associated SNPs. Geographic distribution of the alleles of SNPs that colocalized with inflorescence genes and density distribution of annual precipitation at geographical locations where alleles are distributed for (A) SNP S10_56303321 associated with lower branch length that colocalized the sorghum ortholog of rice *Aberrant Panicle Organization1*, (B) precipitation distribution for accessions carrying each SNP S10_56303321 allele type in the global panel, (C) SNP S3_4750709 associated with lower branch length that colocalized the sorghum ortholog of maize *ramosa2*, (D) precipitation distribution for accessions carrying each SNP S3_4750709 allele type in the global panel, (E) SNP S7_59751994 associated with upper branch length that colocalized the sorghum ortholog of maize *sparse inflorescence1*. The broken lines in density plots represent the mean of each distribution, (F) precipitation distribution for accessions carrying each SNP S7_59751994 allele type in the global panel.

## DISCUSSION

### Genetic architecture of inflorescence adaptation

Nested association mapping can help characterize the genetic architecture of adaptive traits while avoiding some pitfalls of GWAS. This sorghum NAM study provides a high-powered dissection of genetic architecture for global variation in inflorescence morphology, a key trait for adaptation across agroclimatic zones (Harlan and de Wet, 1972; Olatoye *et al.*, 2018). Our study identified many new loci (Table 3) and provided more precise mapping of known loci (Brown *et al.*, 2006; Witt Hmon *et al.*, 2013). Among the known QTL is LBL QTL (qSbLBL7.598), which appears to be pleiotropic with RL (as qSbRL7.598). Previous linkage mapping studies identified association around this same *Dw3* region for QTL associated with rachis length and primary branch length (Brown *et al.*, 2006; Shehzad and Okuno, 2015) and *YUCCA5* was proposed as a candidate gene for the branch length QTL (Brown *et al.*, 2008).

The preponderance of moderate and large effect QTL for four inflorescence morphology traits suggests a predominantly oligogenic trait architecture for inflorescence variation in global sorghum diversity (Supplementary Table 2, Supplementary Figure 3–4, Supplementary File 1-2). Note, a PVE estimate that would be considered “small effect” (e.g. 1%) in a typical biparental study (e.g. 100–300 RILs) is better characterized as “moderate effect: in a NAM population, given the denominator is phenotypic variance across many diverse families and thousands of RILs. In previous studies of sorghum inflorescence in biparental populations, large effect loci (explaining up to 19% of the variance) were found, but effect size of these loci may have been upwardly biased due to the Beavis effect (Xu, 2003). The population size of the NAM (2200 RILs) used in this study should provide a more robust estimation of QTL effect size, which are expected to be accurate with population sizes >1000 (King and Long, 2017). In maize, effect size distribution of loci associated ear and tassel traits has been linked to strong directional selection during maize domestication (Brown *et al.*, 2011; Xue *et al.*, 2016). In sorghum, the moderate to large effect loci identified here may reflect selection towards multiple contrasting fitness optima during the adaptation to contrasting agroclimatic zones, consistent with Fisher’s geometric model under disruptive selection (Orr, 2005; Tenaillon, 2014).

Epistasis may be reflected in asymmetric transgressive variation (Rieseberg *et al.*, 1999; Gaertner *et al.*, 2012). The shift of the RIL means from the mid-parent value in some families, and some strongly skewed trait distributions in NAM RILs, suggest that epistasis may be pervasive (e.g. UBL in SC283 family or RD in SC1103 family; Figure 1). These results supports previous findings in a small population (Ben-Israel *et al.*, 2012). Further evidence for epistatic interactions of additive QTL can be provided by opposite allelic effects of QTL across families (Buckler *et al.*, 2009; Peiffer *et al.*, 2014). Consistent with a hypothesis of gene-by-genetic background epistasis, inflorescence morphology QTL showed opposed allelic effects across families for 63% (82/131) of QTL (Supplementary File 2). Other QTL (16%) identified had consistent allelic effects in all families. These loci may influence inflorescence variation additively across multiple botanical races, or may reflect rare variants in the common parent. Further analyses to map interacting loci will be needed to characterize the role of epistasis in sorghum inflorescence variation (Chen *et al.*, 2018).

Genetic correlation among traits due to linkage or pleiotropy can limit or promote adaptive evolution (Lynch and Walsh, 1998). LBL and RL had high phenotypic correlation (*r* = 0.71, *P*-value < 0.001) and had two major effect QTL that were in common (qSbLBL7.598/qSbRL7.598 and qSbLBL10.563/qSbRL10.563) (Figure 3). Given the large size of NAM population, and concomitant high mapping resolution, if these QTL colocalizations are not due to pleiotropy then linkage must be very tight (e.g. <2 cM). In maize, mutations in the YUCCA-family gene *sparse inflorescence1* led to drastic reduction in both inflorescence rachis length and branch length (Gallavotti *et al.*, 2008), suggesting pleiotropy as a parsimonious explanation for the genetic correlation of these traits. By contrast, the two branch length traits (LBL and UBL) had low phenotypic correlation and lack of colocalization between QTL, suggesting that they are largely under independent genetic control. Studies of the underlying molecular network (e.g. mutant analysis, spatiotemporal expression dynamics) should provide further insight on the basis of pleiotropic versus independent genetic control (Eveland *et al.*, 2014).

Studies have shown evidence of sorghum traits are subject to local adaptation across agroclimatic zones (Morris et al. 2013; Olatoye et al. 2018; Wang et al. 2019). In this study, our results showed independent global geographic distribution of the alleles of inflorescence QTL markers that colocalized with *a priori* genes regulating inflorescence branch traits like lower branch length and upper branch length (Figure 5). This supports previous finding that there is an independent spread of multiple alleles influencing inflorescence traits (Morris et al. 2013). Furthermore, our results showed that these NAM inflorescence QTL markers’ alleles were not strongly associated with annual mean precipitation across global precipitation zones (Figure 5). However, there is evidence for clinal adaptation of sorghum inflorescence to regional precipitation gradient (Olatoye et al. 2018). This suggests that variation at some of these selected genes may not underlie clinal variation of inflorescence to precipitation gradient.

Our comparison of NAM and GWAS QTL highlights that naive GWAS models (GLM) can contain valuable associations signals for adaptive traits. The number of *a priori* candidate genes that colocalized with NAM versus GLM overlaps were higher than *a priori* candidate genes that colocalized with NAM versus CMLM overlaps. This finding suggests that while nominal GLM *P*-values are often inflated, the top associations in simple GLM may reflect true QTL that are not identified in MLM because they are colinear with polygenic variance (Figure 4 and Supplementary File 1) (Huang *et al.*, 2010).

### Evidence of variation in ancestral regulatory networks

Conserved regulatory networks underlying inflorescence development have been identified by comparative mutant and QTL studies (Kellogg, 2007; Zhang and Yuan, 2014). However, it is not yet known whether variation in these ancestral regulatory networks underlies local adaptation of inflorescence morphology. The enrichment of sorghum homologs of grass inflorescence genes at inflorescence QTL suggests that a substantial proportion of sorghum inflorescence variation is due to polymorphism in ancestral regulatory networks. Some of the *a priori* candidate genes that colocalized with inflorescence QTL were sorghum homologs of hormone transporters or biosynthesis enzymes that regulate inflorescence development. One example is at qSbLBL7.598/qSbRL7.598, which was centered on the intragenic region of *YUCCA5* (putative flavin monooxygenase auxin biosynthesis gene) (Figure 3, Supplementary Figure 3–4). This *YUCCA5* gene is a paralog of maize auxin biosynthesis gene *sparse inflorescence1 (Spi1;* 62% similar to *Spi1)* (Figure 3B). The peak SNP for this QTL is also 70 kb from the canonical sorghum height gene and auxin efflux transporter *Dw3* (Sobic.007G163800) (Figure 3B and 3C).

Several other *a priori* candidate genes under QTL are homologs of transcription factors that regulate gene expression during inflorescence meristem differentiation in cereals. For instance, the top UBL QTL (qSbUBL3.475) colocalized with the sorghum ortholog of maize *ramosa2 (ra2)* encoding a C2H2 zinc-finger transcription factor (Sobic.003G052900, 92.7% similarity to *ra2).* In maize and sorghum, the *ra2* transcript is expressed in a group of cells that predicts the position of axillary meristem formation in inflorescence (Bortiri *et al.*, 2006; Eveland *et al.*, 2014). The QTL qSbUBL4.585 colocalized with a sorghum paralog (Sobic.004G237300, 12% similar to *BAD1)* of maize *Branch Angle Defective1 (BAD1)*, a TCP transcription factor that controls cell proliferation in tassel development (Bai *et al.*, 2012).

An LBL and RL QTL (qSbLBL10.563/qSbRL10.563) colocalized with the sorghum ortholog of rice *Aberrant Panicle Organization1 (APO1)* (Sobic.010G220400, 90% similar to *APO1)*, which encodes an F-box protein that regulates inflorescence meristem fate (Figure 3B and 3C) (Ikeda *et al.*, 2007). Sorghum *APO1* was also tagged (inside the gene) by a top branch length-associated SNP in a previous GWAS using the SAP (Morris *et al.*, 2013), strongly suggesting this gene underlies variation for inflorescence compactness. Another RL QTL (qSbRL1.207) colocalized with the sorghum ortholog of rice *TAW1 (TAWAWA1)* gene (Sobic.001G219400, 70% similar to *TAW1).* TAW1 regulates rice inflorescence meristem development (Yoshida *et al.*, 2013), so our study suggests it may condition natural variation for inflorescence morphology in grasses more generally.

### Prospects for genomic prediction of inflorescence morphology

Obtaining locally-adaptive inflorescence morphology is an essential requirement for sorghum breeding programs globally. In field-based phenotypic selection, inflorescence morphology is directly observable prior to pollination. However, a shift to rapid-cycling genomics-enabled breeding in controlled conditions (Watson *et al.*, 2018) would require accurate marker selection or genome prediction of inflorescence morphology along with other agronomic traits. Since the NAM founders originated from diverse agroclimatic zones, genotype-phenotype map we developed should be relevant for sorghum breeding programs globally. More diversity in inflorescence morphology is likely yet to be discovered in sorghum, since ~30% of global variation was not captured in the 11 NAM founder parents (Bouchet *et al.*, 2017). Therefore, increasing the number of families in the NAM population should be beneficial for both increased mapping resolution and allelic diversity. Although this may increase phenotyping burden, the use of high-throughput phenotyping platforms could overcome this challenge (Crowell *et al.*, 2016).

## ACKNOWLEDGEMENTS

The study was carried out using the Beocat High-Performance Computing facility and Integrated Genomics Facility at Kansas State University. This study is contribution no. *[add after acceptance]* from the Kansas Agricultural Experiment Station.

## SUPPLEMENTARY FIGURES AND TABLES

**Table S1:**
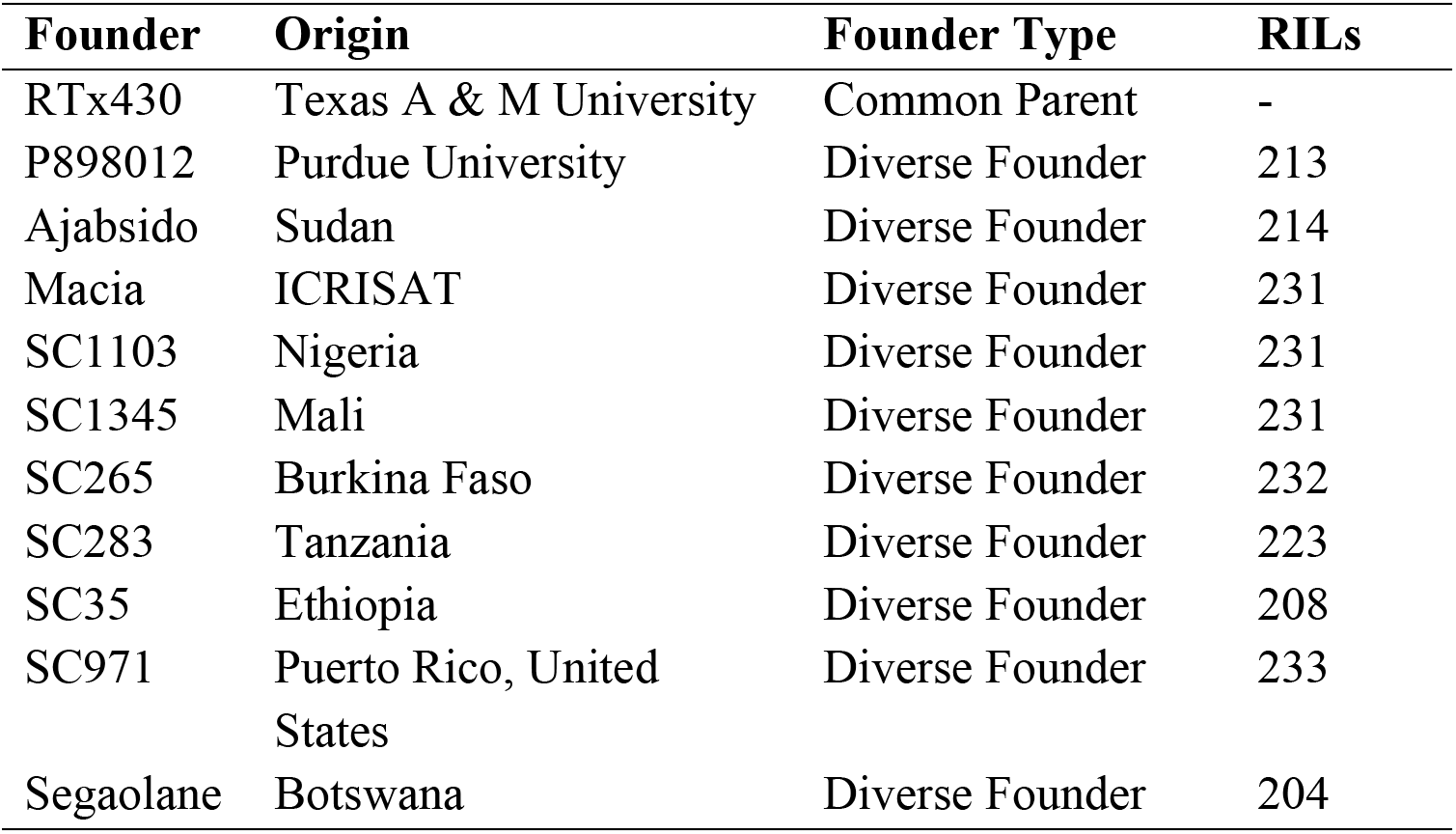
The sorghum NAM founders, their origin, and number of RILs used for each family.

**Table S2:** Within-family and across population additive effect size (AES) for QTL identified using joint linkage mapping with marker nested within family (NJL) and joint linkage with no nesting (JL).

| Trait | Model | Number of QTL | Trait Mean (mm) | Range of WF-AES (mm) | Range of AP-AES (mm) | Range of PVE (%) |
| --- | --- | --- | --- | --- | --- | --- |
| LBL | NJL | 14 | 85 | -30–16 | -4–2 | 0.1–2.0 |
| LBL | JL | 21 | - | -26–19 | -12–8 | 0.6–5.0 |
| UBL | NJL | 1 | 49 | - 29.0–0 | -4 | 3.0 |
| UBL | JL | 17 | - | -33–5 | -11–2 | 0.6–4.0 |
| RL | NJL | 16 | 276 | -44–52 | -10–12 | 0.1–3.0 |
| RL | JL | 22 | - | -49.5–52 | -20–20 | 0.6–3.0 |
| RD | NJL | 9 | 8 | -2.0–1 | -0.4–0.2 | 0.1–1.0 |
| RD | JL | 21 | - | -2.4–1 | -0.9–1.5 | 0.2–1.0 |

<sup>a</sup> Lower branch length (LBL), upper branch length (UBL), rachis length (RL), and rachis diameter (RD).
<sup>b</sup> Quantitative trait loci (QTL).
<sup>c</sup> Within Family Additive Effect Size (WF-AES), and Across Population Additive Effect Size (AP-AES).
<sup>d</sup> Phenotypic Variation Explained (PVE).

**Figure S1:**
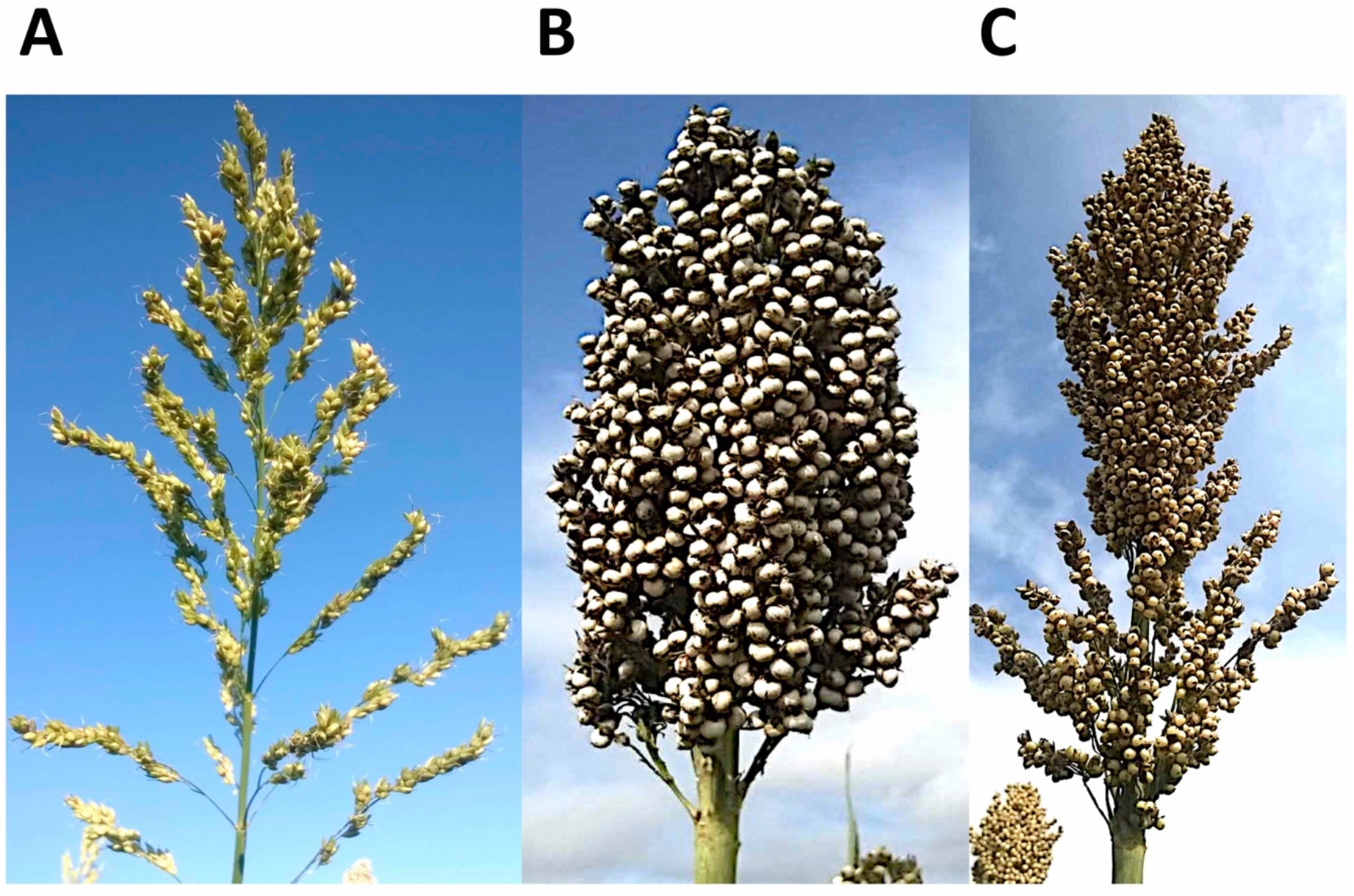
Examples of variation in sorghum inflorescence morphology. (A) Open inflorescence morphology represented by SC1103 parent, (B) compact inflorescence morphology represented by Ajabsido parent, and (C) semi-compact inflorescence morphology as represented by RTx430 the common parent.

**Figure S2:**
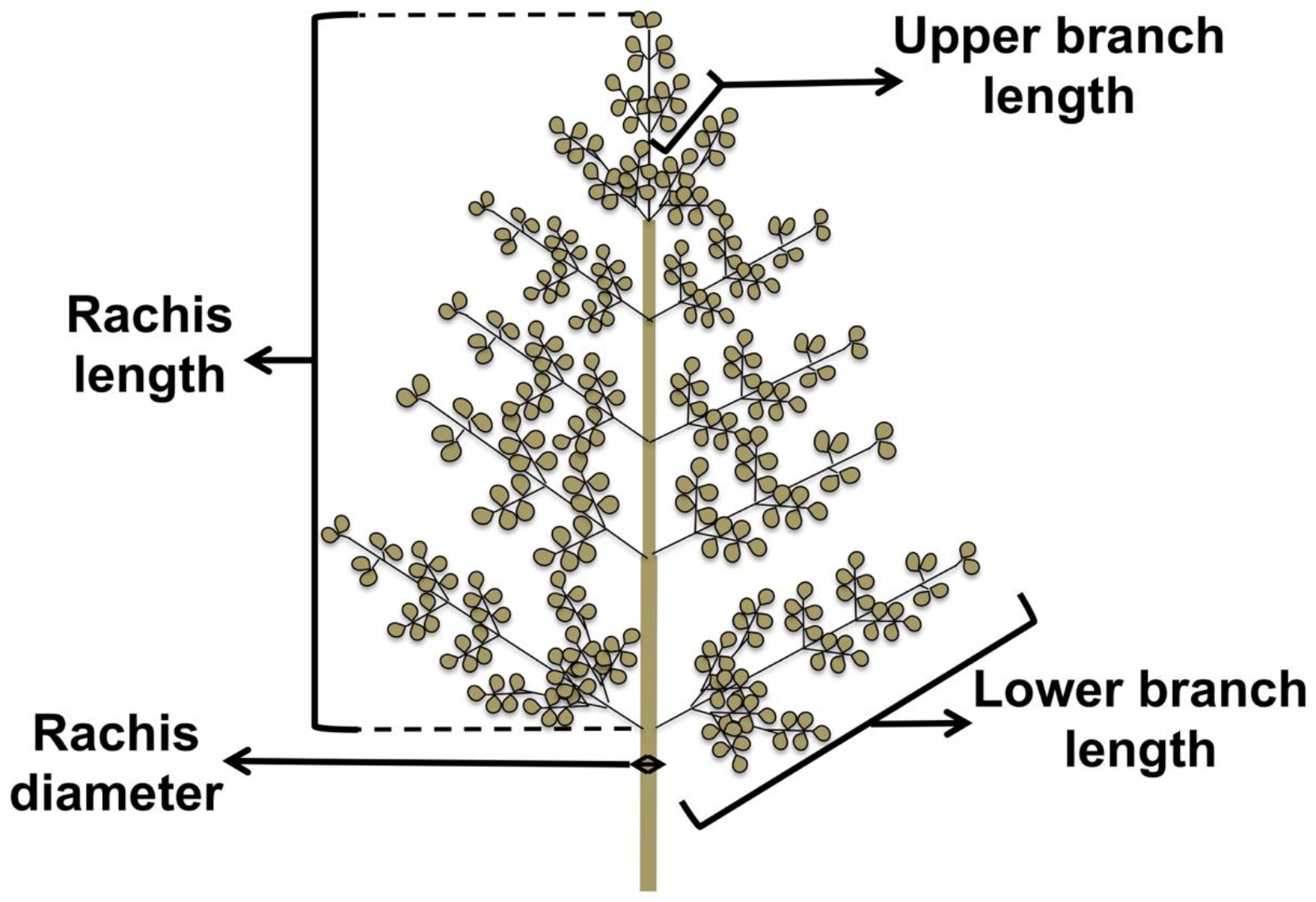
Sorghum inflorescence morphology traits.

**Figure S3:**
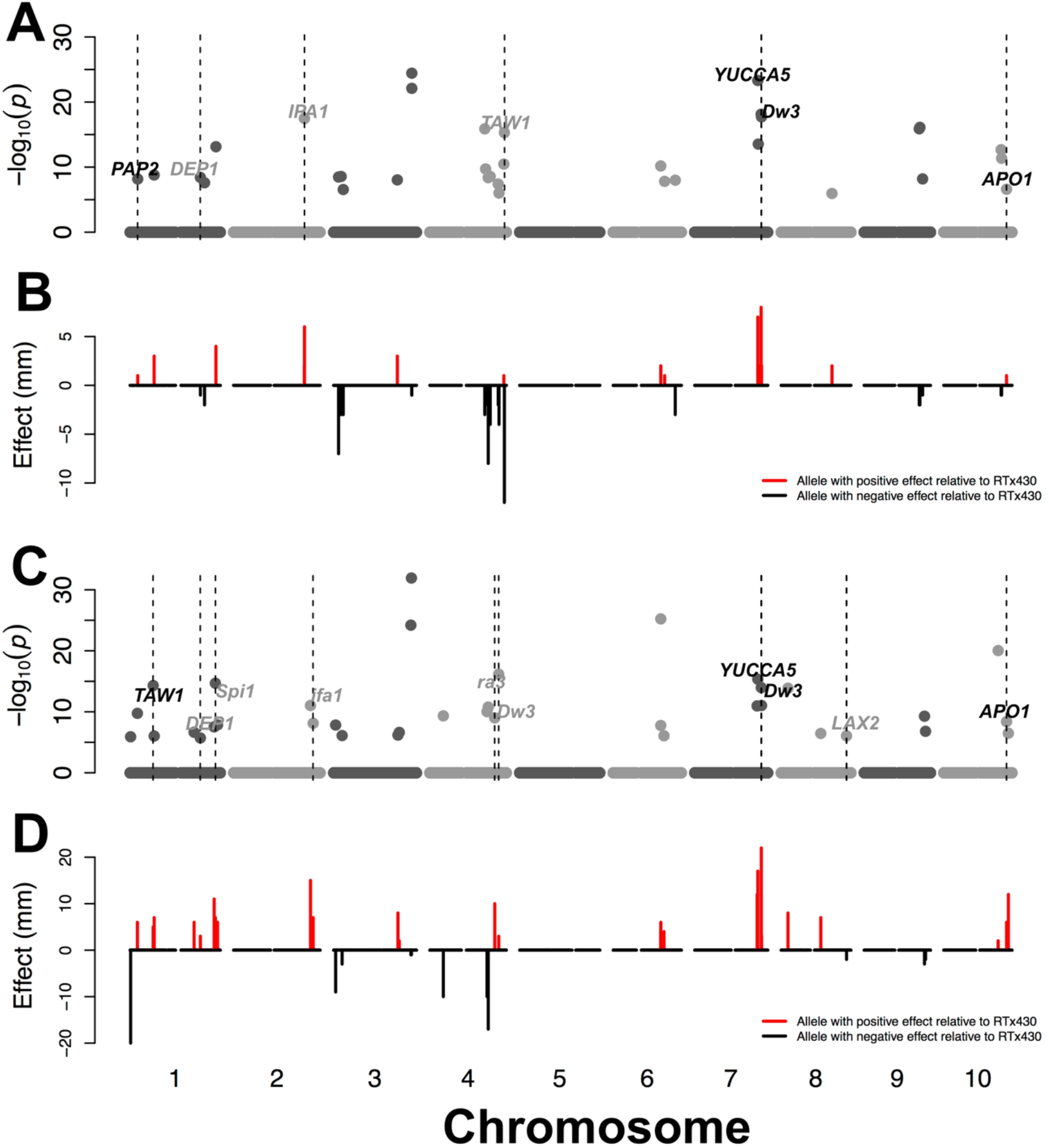
Details of joint linkage QTL mapping for lower branch length and rachis length. Genome positions of loci and loci additive effect associated with lower branch length (A and B) and rachis length (C and D). *A priori* candidate genes that colocalize with QTL within 150 kb are noted. Black text indicates putative sorghum orthologs of *a priori* candidate genes while gray text indicates paralogs.

**Figure S4:**
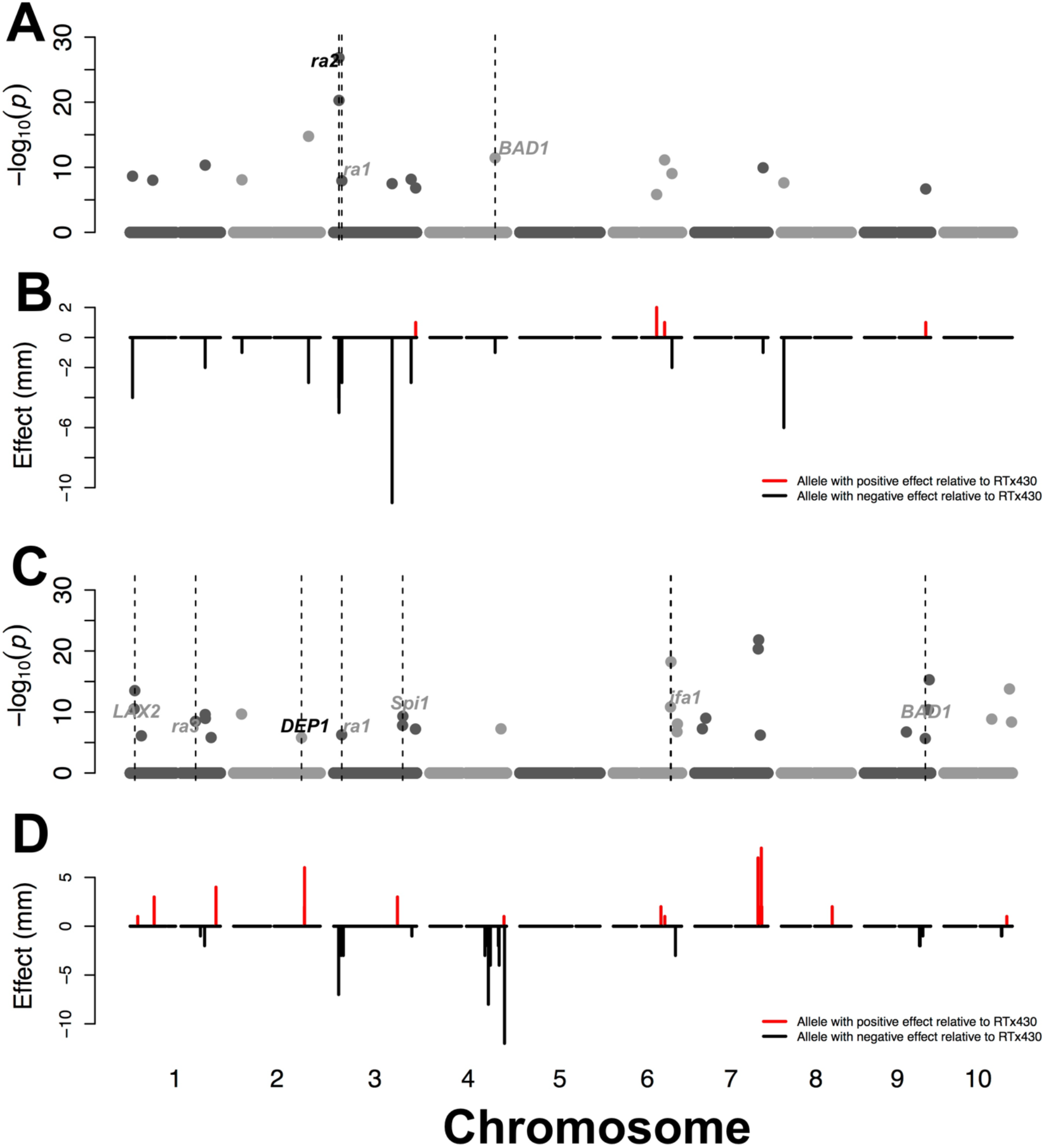
Details of joint linkage QTL mapping for upper branch length and rachis diameter. Genome positions of loci and loci additive effect for upper branch length (A and B) and rachis diameter (C and D). *A priori* candidate genes that colocalize with QTL within 150 kb are noted. Black text indicates putative sorghum orthologs of *a priori* candidate genes while gray text indicates paralogs.

## Literature Cited

Bai F, Reinheimer R, Durantini D, Kellogg EA, Schmidt RJ (2012). TCP transcription factor, BRANCH ANGLE DEFECTIVE 1 (BAD1), is required for normal tassel branch angle formation in maize. Proc Natl Acad Sci.

Bajgain P, Rouse MN, Tsilo TJ, Macharia GK, Bhavani S, Jin Y, et al. (2016). Nested association mapping of stem rust resistance in wheat using genotyping by sequencing. PLOS ONE 11: e0155760.

Barazesh S, McSteen P (2008). Hormonal control of grass inflorescence development. Trends Plant Sci 13: 656–662.

Barrett RDH, Hoekstra HE (2011). Molecular spandrels: tests of adaptation at the genetic level. Nat Rev Genet 12: 767–780.

Bates D, Maechler M, Bolker B, Walker S, Christensen RHB, Singmann H, et al. (2014). lme4: Linear mixed-effects models using Eigen and S4.

Ben-Israel I, Kilian B, Nida H, Fridman E (2012). Heterotic trait locus (HTL) mapping identifies intra-locus interactions that underlie reproductive hybrid vigor in Sorghum bicolor. PLOS ONE 7: e38993.

Bergelson J, Roux F (2010). Towards identifying genes underlying ecologically relevant traits in Arabidopsis thaliana. Nat Rev Genet 11: 867–879.

Bernardo R (2008). Molecular markers and selection for complex traits in plants: Learning from the last 20 years. Crop Sci 48: 1649.

Bortiri E, Chuck G, Vollbrecht E, Rocheford T, Martienssen R, Hake S (2006). Ramosa2 encodes a LATERAL ORGAN BOUNDARY domain protein that determines the fate of stem cells in branch meristems of maize. Plant Cell 18: 574–585.

Bouchet S, Olatoye MO, Marla SR, Perumal R, Tesso T, Yu J, et al. (2017). Increased power to dissect adaptive traits in global sorghum diversity using a nested association mapping population. Genetics 206: 573–585.

Brachi B, Morris GP, Borevitz JO (2011). Genome-wide association studies in plants: the missing heritability is in the field. Genome Biol 12: 232.

Brown P, Klein P, Bortiri E, Acharya C, Rooney W, Kresovich S (2006). Inheritance of inflorescence architecture in sorghum. Theor Appl Genet 113: 931–942.

Brown PJ, Rooney WL, Franks C, Kresovich S (2008). Efficient mapping of plant height quantitative trait loci in a sorghum association population with introgressed dwarfing genes. Genetics 180: 629–637.

Brown PJ, Upadyayula N, Mahone GS, Tian F, Bradbury PJ, Myles S, et al. (2011). Distinct genetic architectures for male and female inflorescence traits of maize. PLoS Genet 7: e1002383.

Buckler ES, Holland JB, Bradbury PJ, Acharya CB, Brown PJ, Browne C, et al. (2009). The genetic architecture of maize flowering time. Science 325: 714–718.

Casa AM, Pressoir G, Brown PJ, Mitchell SE, Rooney WL, Tuinstra MR, et al. (2008). Community resources and strategies for association mapping in sorghum. Crop Sci 48: 30–40.

Chen AH, Ge W, Metcalf W, Jakobsson E, Mainzer LS, Lipka AE (2018). An assessment of true and false positive detection rates of stepwise epistatic model selection as a function of sample size and number of markers. Heredity: 1.

Cooper M, Gho C, Leafgren R, Tang T, Messina C (2014). Breeding drought-tolerant maize hybrids for the US corn-belt: discovery to product. J Exp Bot: eru064.

Crowell S, Korniliev P, Falcão A, Ismail A, Gregorio G, Mezey J, et al. (2016). Genome-wide association and high-resolution phenotyping link Oryza sativa panicle traits to numerous trait-specific QTL clusters. Nat Commun 7: 10527.

De Wet JMJ, Harlan JR, Kurmarohita B (1972). Origin and evolution of guinea sorghums. East Afr Agric For J 38: 114–119.

Eveland AL, Goldshmidt A, Pautler M, Morohashi K, Liseron-Monfils C, Lewis MW, et al. (2014). Regulatory modules controlling maize inflorescence architecture. Genome Res 24:431–443.

Gaertner BE, Parmenter MD, Rockman MV, Kruglyak L, Phillips PC (2012). More than the sum of its parts: a complex epistatic network underlies natural variation in thermal preference behavior in Caenorhabditis elegans. Genetics 192: 1533–1542.

Gallavotti A, Barazesh S, Malcomber S, Hall D, Jackson D, Schmidt RJ, et al. (2008). sparse inflorescence1 encodes a monocot-specific YUCCA-like gene required for vegetative and reproductive development in maize. Proc Natl Acad Sci 105: 15196–15201.

Glaubitz JC, Casstevens TM, Lu F, Harriman J, Elshire RJ, Sun Q, et al. (2014). TASSEL-GBS: A high capacity genotyping by sequencing analysis pipeline. PLoS ONE 9: e90346.

Goodstein DM, Shu S, Howson R, Neupane R, Hayes RD, Fazo J, et al. (2012). Phytozome: a comparative platform for green plant genomics. Nucleic Acids Res 40: D1178–D1186.

Hansen TF (2006). The evolution of genetic architecture. Annu Rev Ecol Evol Syst 37: 123–157.

Harlan JR, de Wet JMJ (1972). A simplified classification of cultivated sorghum. Crop Sci 12: 172–176.

Hermann K, Kuhlemeier C (2011). The genetic architecture of natural variation in flower morphology. Curr Opin Plant Biol 14: 60–65.

Holland JB (2007). Genetic architecture of complex traits in plants. Curr Opin Plant Biol 10: 156–161.

Hu Z, Olatoye MO, Marla S, Morris GP (2019). An integrated genotyping-by-sequencing polymorphism map for over 10,000 sorghum genotypes. Plant Genome 12.

Huang P, Jiang H, Zhu C, Barry K, Jenkins J, Sandor L, et al. (2017). Sparse panicle1 is required for inflorescence development in Setaria viridis and maize. Nat Plants 3: 17054.

Huang X, Wei X, Sang T, Zhao Q, Feng Q, Zhao Y, et al. (2010). Genome-wide association studies of 14 agronomic traits in rice landraces. Nat Genet 42: 961–967.

Hung H-Y, Browne C, Guill K, Coles N, Eller M, Garcia A, et al. (2012). The relationship between parental genetic or phenotypic divergence and progeny variation in the maize nested association mapping population. Heredity 108: 490–499.

Ikeda K, Ito M, Nagasawa N, Kyozuka J, Nagato Y (2007). Rice ABERRANT PANICLE ORGANIZATION 1, encoding an F-box protein, regulates meristem fate. Plant J 51: 1030–1040.

Kellogg EA (2007). Floral displays: genetic control of grass inflorescences. Curr Opin Plant Biol 10: 26–31.

King EG, Long AD (2017). The Beavis effect in next-generation mapping panels in Drosophila melanogaster. G3 Genes Genomes Genet 7: 1643–1652.

Lasky JR, Upadhyaya HD, Ramu P, Deshpande S, Hash CT, Bonnette J, et al. (2015). Genome-environment associations in sorghum landraces predict adaptive traits. Sci Adv 1: e1400218.

Lipka AE, Tian F, Wang Q, Peiffer J, Li M, Bradbury PJ, et al. (2012). GAPIT: genome association and prediction integrated tool. Bioinformatics 28: 2397–2399.

Lynch M, Walsh B (1998). Genetics and Analysis of Quantitative Traits. Sinauer.

Maurer A, Draba V, Jiang Y, Schnaithmann F, Sharma R, Schumann E, et al. (2015). Modelling the genetic architecture of flowering time control in barley through nested association mapping. BMC Genomics 16: 290.

Morris GP, Ramu P, Deshpande SP, Hash CT, Shah T, Upadhyaya HD, et al. (2013). Population genomic and genome-wide association studies of agroclimatic traits in sorghum. Proc Natl Acad Sci 110: 453–458.

Myles S, Peiffer J, Brown PJ, Ersoz ES, Zhang Z, Costich DE, et al. (2009). Association mapping: critical considerations shift from genotyping to experimental design. Plant Cell 21: 2194–2202.

National Research Council (1996). Lost Crops of Africa: Volume I: Grains. National Academy Press, Washington, D.C.

Olatoye MO, Hu Z, Maina F, Morris GP (2018). Genomic signatures of adaptation to a precipitation gradient in Nigerian sorghum. G3 Genes Genomes Genet 8: 3269–3281.

Orr HA (2005). The genetic theory of adaptation: a brief history. Nat Rev Genet 6: 119–127.

Peiffer JA, Romay MC, Gore MA, Flint-Garcia SA, Zhang Z, Millard MJ, et al. (2014). The genetic architecture of maize height. Genetics 196: 1337–1356.

Poland JA, Bradbury PJ, Buckler ES, Nelson RJ (2011). Genome-wide nested association mapping of quantitative resistance to northern leaf blight in maize. Proc Natl Acad Sci 108: 6893–6898.

Rieseberg LH, Archer MA, Wayne RK (1999). Transgressive segregation, adaptation and speciation. Heredity 83: 363–372.

Segura V, Vilhjálmsson BJ, Platt A, Korte A, Seren Ü, Long Q, et al. (2012). An efficient multilocus mixed-model approach for genome-wide association studies in structured populations. Nat Genet 44: 825–830.

Shehzad T, Okuno K (2015). QTL mapping for yield and yield-contributing traits in sorghum (Sorghum bicolor (L.) Moench) with genome-based SSR markers. Euphytica 203: 17–31.

Tanaka W, Pautler M, Jackson D, Hirano HY (2013). Grass meristems II: Inflorescence architecture, flower development and meristem fate. Plant Cell Physiol 54: 313–324.

Tenaillon O (2014). The utility of Fisher’s geometric model in evolutionary genetics. Annu Rev Ecol Evol Syst 45: 179–201.

Timpson NJ, Greenwood CMT, Soranzo N, Lawson DJ, Richards JB (2018). Genetic architecture: the shape of the genetic contribution to human traits and disease. Nat Rev Genet 19: 110–124.

Utz HF, Melchinger AE, Schön CC (2000). Bias and sampling error of the estimated proportion of genotypic variance explained by quantitative trait loci determined from experimental data in maize using cross validation and validation with independent samples. Genetics 154:1839–1849.

Watson A, Ghosh S, Williams MJ, Cuddy WS, Simmonds J, Rey M-D, et al. (2018). Speed breeding is a powerful tool to accelerate crop research and breeding. Nat Plants 4: 23.

Witt Hmon KP, Shehzad T, Okuno K (2013). Variation in inflorescence architecture associated with yield components in a sorghum germplasm. Plant Genet Resour 11: 258–265.

Wu X, Li Y, Shi Y, Song Y, Zhang D, Li C, et al. (2016). Joint-linkage mapping and GWAS reveal extensive genetic loci that regulate male inflorescence size in maize. Plant Biotechnol J 14: 1551–1562.

Würschum T, Liu W, Gowda M, Maurer HP, Fischer S, Schechert A, et al. (2012). Comparison of biometrical models for joint linkage association mapping. Heredity 108: 332–340.

Xu S (2003). Theoretical basis of the Beavis Effect. Genetics 165: 2259–2268.

Xue S, Bradbury PJ, Casstevens T, Holland JB (2016). Genetic architecture of domestication-related traits in maize. Genetics 204: 99–113.

Yoshida A, Sasao M, Yasuno N, Takagi K, Daimon Y, Chen R, et al. (2013). TAWAWA1, a regulator of rice inflorescence architecture, functions through the suppression of meristem phase transition. Proc Natl Acad Sci U S A 110: 767–772.

Yu J, Holland JB, McMullen MD, Buckler ES (2008). Genetic design and statistical power of nested association mapping in maize. Genetics 178: 539–551.

Yu J, Pressoir G, Briggs WH, Bi IV, Yamasaki M, Doebley JF, et al. (2006). A unified mixed-model method for association mapping that accounts for multiple levels of relatedness. Nat Genet 38: 203–208.

Zhang D, Yuan Z (2014). Molecular control of grass inflorescence development. Annu Rev Plant Biol 65: 553–578.

